# Temporal Optimization of Radiation Therapy to Heterogeneous Tumour Populations and Cancer Stem Cells

**DOI:** 10.1101/2022.01.26.477741

**Authors:** Cameron Meaney, Mohammad Kohandel, Arian Novruzi

## Abstract

External beam radiation therapy is a key part of modern cancer treatments which uses high doses of radiation to destroy tumour cells. Despite its widespread usage and extensive study in theoretical, experimental, and clinical works, many questions still remain about how best to administer it. Many mathematical studies have examined optimal scheduling of radiotherapy, and most come to similar conclusions. Importantly though, these studies generally assume intratumoral homogeneity. But in recent years, it has become clear that tumours are not homogeneous masses of cancerous cells, but wildly heterogeneous masses with various subpopulations which grow and respond to treatment differently. One subpopulation of particular importance is cancer stem cells (CSCs) which are known to exhibit higher radioresistence compared with non-CSCs. Knowledge of these differences between cell types could theoretically lead to changes in optimal treatment scheduling. Only a few studies have examined this question, and interestingly, they arrive at apparent conflicting results. However, an understanding of their assumptions reveals a key difference which leads to their differing conclusions.

In this paper, we generalize the problem of temporal optimization of dose distribution of radiation therapy to a two cell type model. We do so by creating a mathematical model and a numerical optimization algorithm to find the distribution of dose which leads to optimal cell kill. We then create a data set of optimization solutions and use data analysis tools to learn the relationships between model parameters and the qualitative behaviour of optimization results. Analysis of the model and discussion of biological importance are provided throughout. We find that the key factor in predicting the behaviour of the optimal distribution of radiation is the ratio between the radiosensitivities of the present cell types. These results can provide guidance for treatment in cases where clinicians have knowledge of tumour heterogeneity and of the abundance of CSCs.

## 1 Introduction

As one of the pillar of modern cancer treatment, radiation therapy has been the subject of countless research works. Despite its origins in cancer medicine tracing back to over a century ago, many questions and challenges still remain. Of particular interest in recent years, improving our ability to accurately predict the efficacy of a particular course of radiation prior to treatment remains an important clinical goal. In theory, if the efficacy of a given treatment prescription can be predicted prior to administration, then many treatments can be considered and the optimal selected. Furthermore, if patient-specific knowledge can be incorporated into these predictions, then patient-specific optimization of radiation becomes a reality. Typically, researchers rely on mathematical and computational models to evaluate such predictions, and many promising advancements have been made. In practice however, this has proven far easier said than done, since patient-specific measurements are often difficult and costly to obtain and can come with significant uncertainty. Furthermore, the sheer number of physical and biological factors at play in predicting treatment outcomes makes accurate mathematical predictions challenging.

Many previous works have examined different ways to optimize radiation therapy including temporally [1, 3, 6, 11, 12, 16, 25], spatially [2, 14, 18, 23], and patient scheduling [10]. Generally, these studies perform mathematical optimizations, occurring in four steps: first, select metrics; second, determine constraints; third, write governing model of treatment response; and fourth, perform mathematical optimization. When discussing radiation, several different metrics can be considered. The most common is to maximize cell kill, but others such as maximizing tumour control probability (TCP), minimizing administered dose, and minimizing tissue complication probability are also commonly considered. Some studies consider multiple metrics, typically by forming a combined metric with weight coefficients. Constraints for the mathematical problem are chosen based on the physical and biological limitations of the situation. In some cases, this means limiting the total administered dose to a prescribed amount over the course of a day or a full treatment period. In other cases, constraints include timing considerations such as prohibiting radiation given on weekends. Models of treatment response typically involve ordinary or partial differential equations (ODEs or PDEs) and use established models of radiation effect such as the linear-quadratic (LQ) model. These models determine the effect of a chosen candidate treatment with respect to the chosen metric. Once the metric, constraints, and model have been fixed, a mathematical optimization can be performed where an optimal is selected from the many candidate treatments.

In this paper, we consider the problem of finding the temporal distribution of radiation dose which minimizes the total remaining number of cells at the end of treatment under a set of clinically-relevant constraints. Several previous works have examined similar questions. In Wein et al [25], the authors considered a spatially-varying population of homogeneous tumour cells and used dynamic reprogrammming to find the radiation schedule that optimizes the TCP under constraints to the biologically effective dose (BED). They found that allowing the fractional dose amount to change over the course of treatment allowed for a higher TCP than the clinically-standard constant dose rate. Altman et al [3] obtained a similar result, showing that when maximizing cell kill over a homogeneous population of tumour cells subject to constrained total dose, the optimal distribution of radiation was again when the dose rate was allowed to vary throughout treatment. In Kim et al [14], the authors considered maximizing the BED while constraining the dose for nearby organs-at-risk (OARs). They argued that due to recent technological advancements allowing more precise localization of radiation dose on a tumour core, treatment optimizations can focus primarily on maximizing anticancer effect. Even when including spatial effects in their optimization, the authors found that varying the dose per fraction could lead to an improvement over uniform schedules. Some analytical results were derived in Mizuta et al [19] related to optimal temporal dose distributions where cell kill was to be optimized and BED to an OAR minimized. They provided a theorem stating that in cases without tumour repolulation, if the OAR is sufficiently sensitive, then the optimal dose distribution is uniform over the treatment length, but if the OAR is not sufficiently sensitive, then the optimal distribution is to focus the allowable radiation into the least number of fractions possible. Bortfeld et al [6] built off of the work of Mizuta et al by including the effect of tumour repopulation. They similarly showed that in cases where the OAR is less sensitive, the optimal distribution was to focus radiation into the smallest number of fractions possible. Through their analysis, they were further able to conclude that for tumours with faster growth rates, the optimal distribution of radiation tended to be spread over a shorter time.

Despite the important differences between the above works with respect their particular problem formulation, they all arrive at similar conclusions. Specifically, that a nonuniform fractional dose distribution can outperform the clinically-standard uniform distribution, and that the optimal case tends to involve focusing allowable dose over shorter periods of time at higher dose rates. Clearly, such results are of tremendous clinical importance. However, the applicability of these works is somewhat lessened by the omission of a key feature of tumours: cellular heterogeneity. Each of the above works considers homogeneous tumours in which all cells proliferate, invade, and respond to treatment in the same way. In reality, tumours are heterogeneous and contain many different types of cells which can act in different ways. Of particular importance, cancer stem cells (CSCs) are a type of tumour cell capable of self-renewal and differentiation. They are thought to represent a small subpopulation of the full tumour, yet nevertheless be the primary drivers of tumour growth and repopulation. Countless recent studies have noted their importance in understanding cancer progression and treatment ([4, 5], for example). Studies have also noted experimental techniques which could be used to identify stem-like subpopulations, relying typically on protein biomarkers [20]. Despite the imperfect natures of these biomarker methods, differences between biomarker positive and biomarker negative cells can be observed. With respect to radiation therapy, studies have examined the difference in response to radiation by CSCs and non-CSCs [15, 21] (or biomarker +*/*− cells), finding that CSCs are commonly less sensitive to radiotherapy than their non-CSC counterparts. Furthermore, they observe that post-radiation tumours tend to have become less heterogeneous as sensitive cells are removed by radiation and all that remains are the resistant stem cells. Such observations suggest that considering a heterogeneous tumour population may be a crucial feature in developing accurate models of radiation optimization. Indeed, many recent works have done just this.

Leder et al [16] considered a mathematical model with two distinct cell types: a radioresistant CSC-like cell and a radiosensitive non-CSC-like cell. Using a two compartment mathematical model along with a Monte Carlo optimization method, they found that mathematically optimized radiation schedules led to a longer survival time and greater reduction in tumour volume than the standard schedule in glioblastoma-bearing mice. Notably, their optimal schedule no longer had the qualitative behaviour observed in the one cell type cases described above. In fact, the hypofractionated (less fractions at higher dose rates) case performed the worst out of the four tested. In Forouzannia et al [11], the authors also considered a two cell type model and included plasticity rates between their CSC and non-CSC compartments. They considered five different schedules of radiation, and sought to find which schedules led to the minimum remaining cells and to find the resulting fraction of CSCs with respect to the full tumour population. Their results were similar to [16] in that neither the standard of care nor hypofractionation was optimal. In contrast however, Galochkina et al [12] also considered a two cell type model but arrived at the opposite result of [11] and [16]. They found that the uniform distribution of dose was typically optimal, and that cases with different optima generally only provided insignificant benefits. However, in [12]’s two cell model, the only difference between the cell types was their replenishment rates - importantly, they were assumed to be affected by radiotherapy in the same way. Given the differing conclusions of these models, it would be reasonable to suspect that a key force in determining the qualitative nature of an optimal radiation schedule is the difference in radiation responses between the cell types as opposed to differences in repopulation or plasticity rates. In this work, we set out to answer this question and determine which situations lead to qualitatively different optimal radiation schedules.

We begin the methods section by detailing the two cell type model that we use to determine the effect of radiation therapy. We then explain our optimization problem, including choice of metric and solution constraints. We provide some important analytical analysis of our model and its possible solutions, specifically proving that solutions to our model is typically ‘bang-bang’ in nature. Finally, we explain the projected gradient descent optimization technique that we employ to find the optimal solution. In the results section, we provide some example solutions of the optimization problem and distinguish between the two different qualitative categories into which solutions fall. We then provide analysis using tree classifiers of the key parameters which determine the behaviour of results, connect this analysis to previous literature, and discuss the clinical relevance. In the conclusion, the work is summarized, and directions for future research are noted.

## 2 Methods

### 2.1 Model

In this work, we consider a tumour consisting of two distinct populations of cells, denoted by *n*_1_ and *n*_2_. We use a system of two ODEs, shown in equations (1) and (2) and in Figure 1, to govern the time evolution of the cell populations.

**Figure 1:**
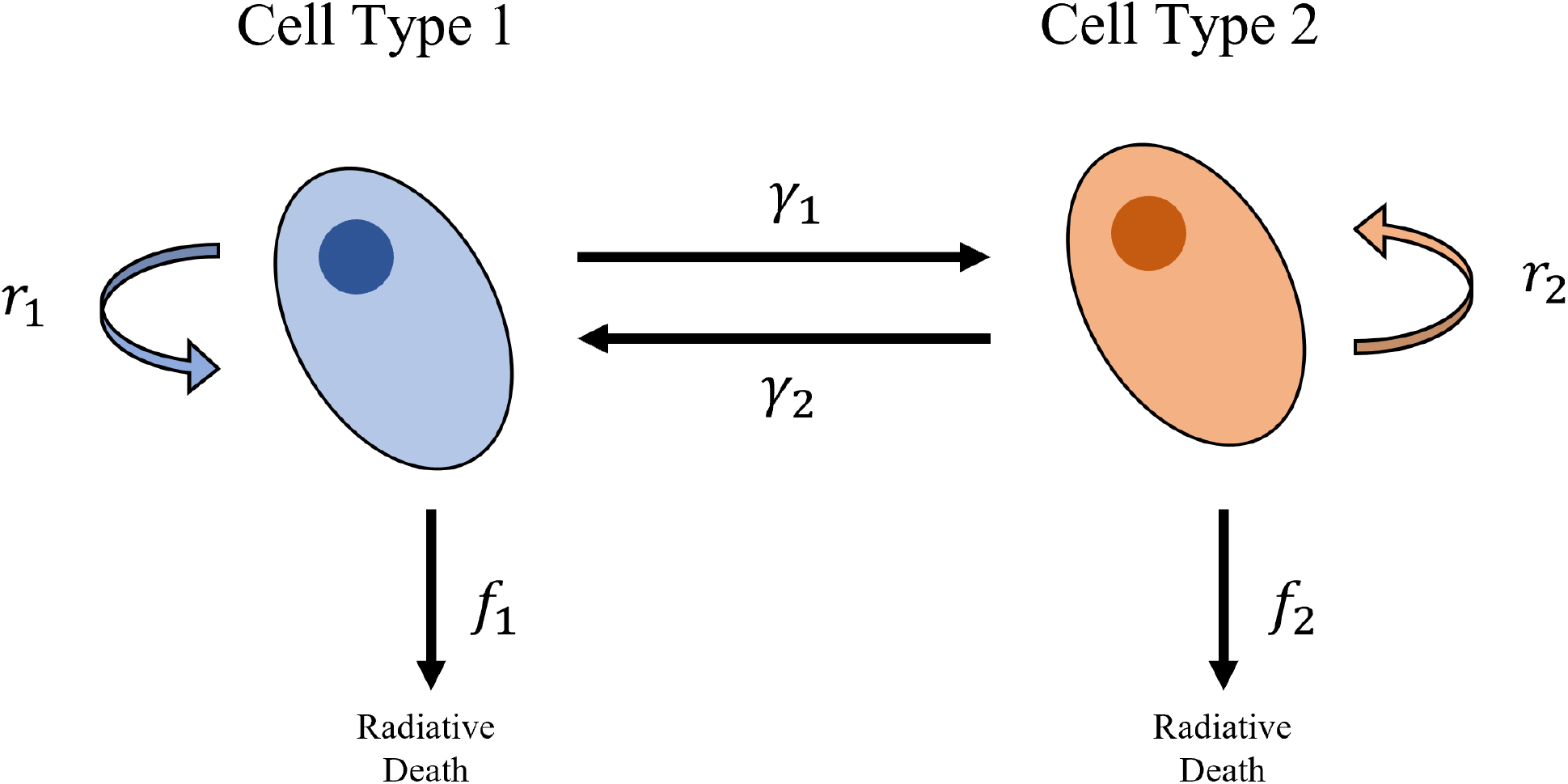
Schematic of the two cell type compartmental ODE model in equations (1) and (2). Each cell type is assumed to proliferate according to a logistic growth law with proliferation rate *r*_*i*_. The plasticity rates between the cells types are given by *γ*_*i*_. Cells are killed by radiation through the functions *f*_*i*_ which depend on the radiobiological parameters *α*_*k*_ and *β*_*k*_.

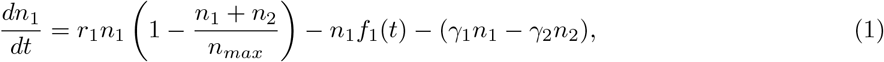

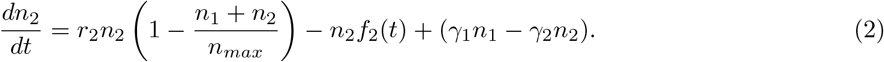

Each cell type is assumed to proliferate according to a combined logistic growth law which is shown in the first term of each equation. The parameter *n*_*max*_ is the cell carrying capacity: the maximum number of cells that the tumour can sustain. As the total number of cells, *n*_1_ + *n*_2_, approaches *n*_*max*_, the rate of increase of the cell population decreases. The proliferation rate parameters *r*_1_ and *r*_2_ differentiate the rates of growth between the cell types, with higher values corresponding to faster growth. When the total cell population is low compared to *n*_*max*_, *r*_1_ and *r*_2_ are equivalent to exponential growth rates. The final term in each equation describes the plasticity between the cell types: cells of type 1 turning into cells of type 2 and the converse. Interestingly, recent works have show the bidirectional nature of plasticity between CSCs and non-CSCs [7, 17, 24]. For this reason, we include bidirectional plasticity in our model with respective plasticity rates of *γ*_1_ (type 1 to type 2) and *γ*_2_ (type 2 to type 1). The middle term of each equation describes the effect of radiotherapy on the cells, which is assumed to be governed by the well-known LQ model and is included in the functions *f*_1_(*t*) and *f*_2_(*t*) in its differential form. In our model, we assume that radiation is applied in a finite number of fractions and denote this number as *N*. We denote the time midpoint of fraction number *i* as *τ*_*i*_, its length by 2*ω*_*i*_, and it dose rate (in Gy/day) by *a*_*i*_. With this formulation, we can write the functions *f*_1_(*t*) and *f*_2_(*t*) as shown in equations (3) and (4).

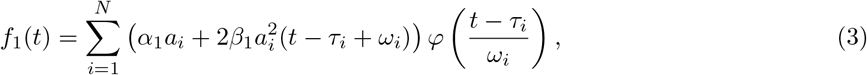

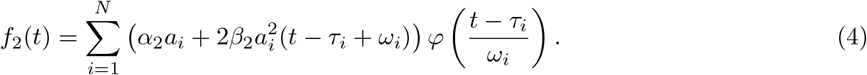

Note that 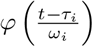 is a rectangular window function centred at time *τ*_*i*_ with width 2*ω*_*i*_ given by

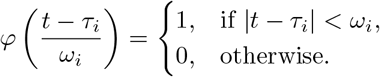

This defines the dose rate as an on-again-off-again function: *a*_*i*_ when radiation is being applied and zero when it’s not. Also note the different values of the radiobiological parameters *α*_*k*_ and *β*_*k*_ for the different cell types. Higher values of these parameters mean that the cell type has a higher sensitivity to radiation. A radiation treatment schedule is therefore fully characterized by the three N-dimensional vectors *a* = (*a*_1_, …, *a*_*N*_) (fraction dose rates), *τ* = (*τ*_1_, …, *τ*_*N*_) (fraction time midpoints), and *ω* = (*ω*_1_, …, *ω*_*N*_) (fraction half lengths). In this work, we are primarily interested in optimizing the distribution of dose over a treatment period, so we make two key assumptions on candidate radiation schedules. First, we assume that the length of each fraction is the same, meaning *ω*_*i*_ = *ω*_*j*_, ∀ *i, j*. Second, we assume that the fraction midpoints are fixed in time, once per day at the same time each day, meaning *τ*_*i*+1_ = *τ*_*i*_ + 1 day, ∀ *i*. Note that this does not mean that radiation is administered every day: if *a*_*i*_ = 0, then no radiation will be given on day *i*. We seek to find the distribution of dose over the possible times, determined by the selection of *a* alone, that leads to the maximum cell kill. For any choice of *a*, the model can be solved to yield a population vs. time plot for each cell type and the total, incorporating the effect of radiation.

As one might expect, without constraints on the applied radiation, the optimal cell kill would result from simply increasing the dose rate of each fraction to its maximal level. But of course, in clinical practice, there are various constraints that prevent this from being a reasonable solution. In our optimization, we consider two constraints on our fractional dose rate schedule: total dose and maximal dose. We first insist that the total dose administered over all fractions is less than or equal to the total allowable amount, and for maximal dose, we simply impose an upper bound on the dose rate given over each fraction, which read as

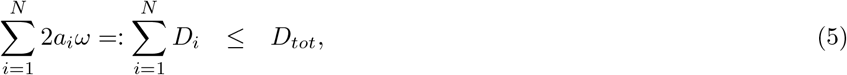

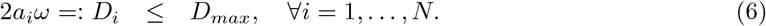

Clinically, the constraint (5) is applied to prevent the patient from receiving dangerous levels of radiation beyond what is necessary for treatment. We expect the inequality in (5) to be saturated in the optimal case. The clinical rationale for (6) is because of the different response times of healthy and cancerous tissues. As healthy tissues generally have a higher capacity to repair DNA damage, dose fractionation can limit the damage to healthy tissues while still incurring high damage to diseased tissues. Without this constraint, the model would be free to converge to a single hit of radiation at a dangerously high dose rate, removing the fractionation altogether. Here, we will exclusively consider cases where *ND*_*max*_ *> D*_*tot*_, meaning that applying the maximal dose per fraction will result in surpassing the limit in total dose as these are the clinically relevant cases and the ones which produce interesting mathematical results. In cases where *ND*_*max*_≤ *D*_*tot*_, the optimal result is trivially to apply *D*_*max*_ at every fraction.

We therefore seek to answer the following question. For a tumour consisting of two distinct subpopulations of cancer cells and a fractionated dose schedule constrained by total dose and maximal dose, what choice of radiation dose rates, *a*, result in the minimal population of total cells remaining after treatment. In our analysis, we generalize to minimizing a weighted sum of cells, *e* = *p*_1_*n*_1_(*T*) +*p*_2_*n*_2_(*T*) for the end of treatment time, *T*. Mathematically, we seek to find

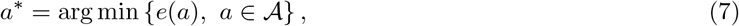

where 𝒜 is the vector space of fractional dose rates *a* = (*a*_1_, …, *a*_*N*_) ∈ ℝ^*N*^ satisfying (5) and (6).

### 2.2 Bang-Bang Structure of the Optimal Radiation Treatment

It is known, see for example [8], that the optimal distribution of resources in population models is of bang-bang type, i.e. the most favorable distribution of resources for a species to survive is when the resources are distributed spatially with patches of maximal or minimal values. If we consider our variable *a* as similar to the growth rate variable *m* in [8], our result represents a time extension of the bang-bang type result of the optimal solution.

In this section, we will show that modulo a subset where the gradient is constant, the solution to (7) is of bang-bang type, i.e. the components of optimal radiation 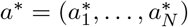 are of three types: i) they are maximal equal to *D*_*max*_, ii) they are minimal equal to 0, or iii) they are such that derivative of the energy *e* on the direction of those components is constant. In our numerical experiments, we always found that the optimal solution *a*\* is purely of bang-bang type, i.e. the set of components of *a*\* where the energy is constant is empty.

Note that the solution of (1)-(2) is understood in the following sense. We say (*n*_1_, *n*_2_) solves (1)-(2) if *n*_1_, *n*_2_ ∈ *W* ^1,∞^(0, *T*) and

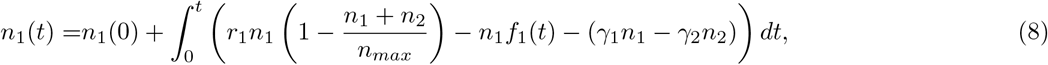

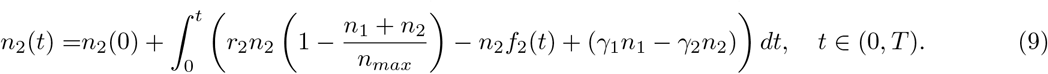

We start with an a priori estimate for the solution (*n*_1_, *n*_2_) of (1)-(2).

#### Proposition 2.1.

*Let F* (*t, n*_1_, *n*_2_) = (*F*_1_(*t, n*_1_, *n*_2_), *F*_2_(*t, n*_1_, *n*_2_)) *with*

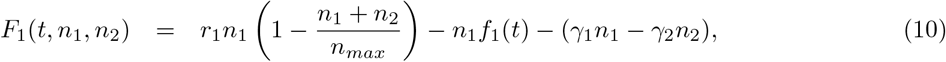

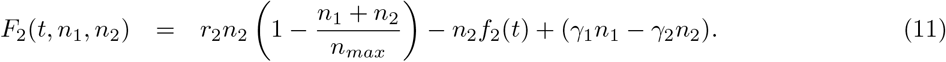

*Assume* (1)*-*(2) *has a solution* (*n*_1_, *n*_2_) ∈ *W* ^1,∞^((0, *T*); ℝ^2^). *If n*_1_(0) ≥ 0, *n*_2_(0) ≥ 0 *then*

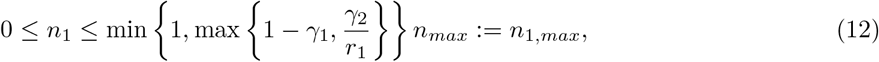

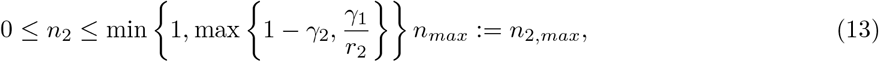

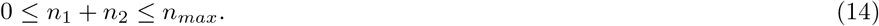

**Proof**. Note that *F*_1_(*t*, 0, *n*_2_) ≥0, *F*_2_(*t, n*_1_, 0) ≥ 0 for all *n*_1_ ≥0, *n*_2_≥ 0. This implies *n*_1_≥ 0, *n*_2_ ≥0, see for example [22, Lemma 1.1]. Furthermore,

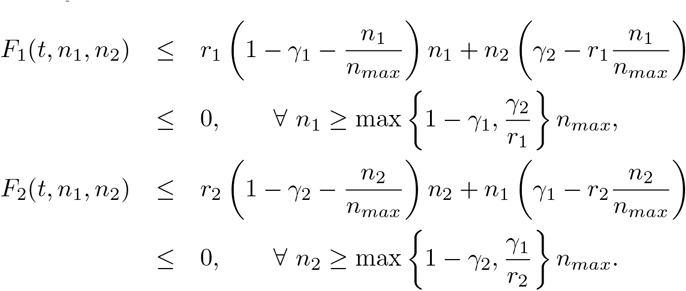

This implies 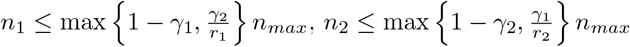. Finally, we have

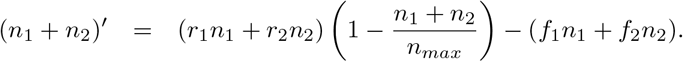

Considering the right hand side of the previous equation as a function of *n*_1_ + *n*_2_, we see that it is negative as soon as *n*_1_ + *n*_2_ = *n*_*max*_, which implies *n*_1_ + *n*_2_ ≤ *n*_*max*_, whence in particular *n*_1_ ≤ *n*_*max*_ and *n*_2_ ≤ *n*_*max*_. □

#### Theorem 2.2.

*The problem* (1)*-*(2) *has a unique solution* (*n*_1_, *n*_2_) ∈ *W* ^1,∞^((0, *T*); ℝ^2^) *for every a* ∈ ℝ^*N*^.

**Proof**. Note that *F* is a Carathéodory function and from [9, Theorem 1.1, Chapter 2], there exists a local in time absolutely continuous solution. Taking into account the form of *F*, which is *C*^∞^ in (*n*_1_, *n*_2_) and *L*^∞^ in *t*, it implies that the local solution is *W* ^1,∞^. As the solution remains bounded, see Proposition 2.1, it extends to *W* ^1,∞^((0, *T*); ℝ^2^) and even to *W* ^1,∞^(ℝ; ℝ^2^).

For the uniqueness we note that for every two solutions *n* = (*n*_1_, *n*_2_), *ñ* = (*ñ*_1_, *ñ*_2_) the function *F* satisfies | *F* (*t, n*_1_, *n*_2_) −*F* (*t, ñ*_1_, *ñ*_1_) |≤ *C*(*n*_1_− *ñ*_1_ | +| *n*_2_− *ñ*_2_ |), because *n* and *ñ* are bounded. Then the uniqueness follows from Gronwall inequality. □

#### Proposition 2.3.

*The functions a* ⟼ *n*_1_(· ; *a*) *and a* ⟼ *n*_2_(· ; *a*) *are C*^∞^ *with respect to a in W* ^1,∞^(0, *T*). *Their first derivative with respect to (wrt) a satisfy*

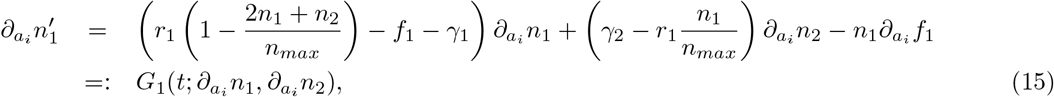

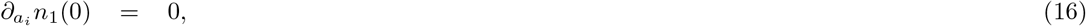

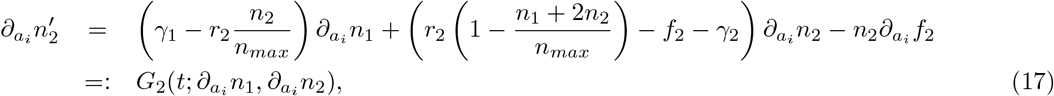

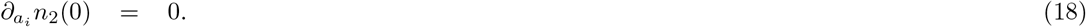

*As a consequence, n*_1_(*T* ; *a*), *n*_2_(*T* ; *a*) *and e*(*a*) *are C*^∞^ *with respect to a and*

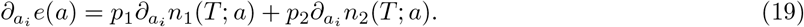

**Proof**. First we show that *n*_1_ and *n*_2_ are *C*^∞^ with respect to *a* in *W* ^1,∞^(0, *T*). This can be done easily by using implicit function theorem as follows. Let *H* = (*H*_1_, *H*_2_) : ℝ^*N*^ × *W* ^1,∞^((0, *T*); ℝ^2^) ⟼ *W* ^1,∞^((0, *T*), ℝ^2^) be given as follows: for *a* ∈ ℝ^*N*^ and *v* = (*v*_1_, *v*_2_) ∈ *W* ^1,∞^((0, *T*); ℝ^2^) we define

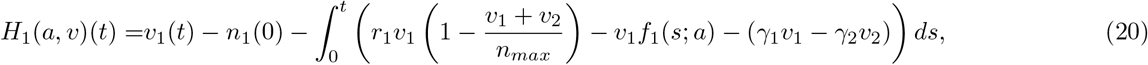

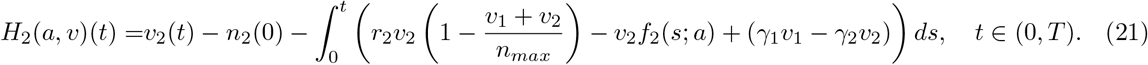

Now let *a*^0^ ∈ ℝ^*N*^ be fixed. We have *H*(*a*^0^, *n*(·; *a*^0^)) = 0 and *H* is *C*^∞^ wrt (*a, v*_1_, *v*_2_). The derivative of *H* wrt to *v* at (*a*^0^; *n*(·; *a*^0^)) in direction *w* is given by

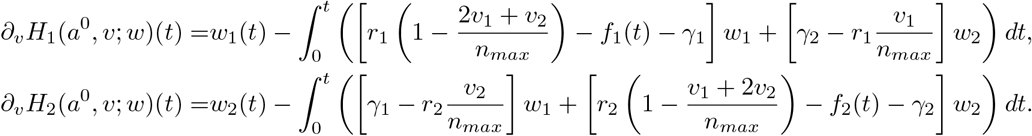

Like for the existence of (*n*_1_, *n*_2_) in Theorem 2.2, it is easy to show that *∂*_*v*_*H*(*a*^0^; *n*(·; *a*^0^)) defines an isomorphism from *W* ^1,∞^((0, *T*); ℝ^2^) to itself. The implicit function theorem implies the existence of a *C*^∞^ map *z* from an open neighbourhood *A*_0_∈ ℝ^*N*^ of *a*^0^ into *W* ^1,∞^((0, *T*); ℝ^2^) such that *H*(*a, z*(·; *a*)) = 0 for all *a* in *A*_0_. The uniqueness of solution for problem (1)-(2) implies *n*(·; *a*) = *z*(·; *a*), which proves the *C*^∞^ differentiability of *n* wrt *a* in *W* ^1,∞^((0, *T*); ℝ^2^).

The *C*^∞^ differentiability wrt *a* of *n* in *W* ^1,∞^((0, *T*); ℝ^2^) and the continuous embedding of *W* ^1,∞^((0, *T*); ℝ^2^) in *C*^0^([0, *T* ]; ℝ^2^) implies also the *C*^∞^ differentiability wrt *a* of *n*(*T* ; *a*) and so of *e*(*a*). Finally, the formulas (15)-(18) and (19) are proved easily from direct calculus. □

#### Theorem 2.4.

*The minimization problem* (7) *has a solution*.

**Proof**. Let (*a*^*k*^), *a*^*k*^, ∈ 𝒜, be a minimization sequence of (7). As 𝒜 is bounded we get that (*a*^*k*^) is bounded in ℝ^*N*^, and from Proposition 2.1 we have 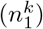 and 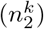, where 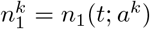 and 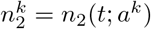, ares bounded in *W* ^1,∞^(0, *T*). It implies:

i. there exists a subsequence of (*a*^*k*^), still denoted (*a*^*k*^), converging to a certain *a*\*, and
ii. there exists a subsequence of 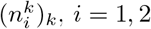, still denoted 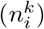, converging in *C*^0^([0, *T* ]) to 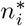.

Clearly *a*\* ∈ 𝒜. By using a standard diagonal procedure, we may assume without loss of generality that 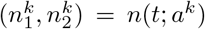. Replacing *a*^*k*^, 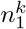 and 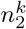 in (8)-(9) and passing in limit shows that 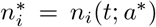, *i* = 1, 2. As *e*(*a*) is continuous wrt *a* it follows that *a*\* solves (7). □

#### Lemma 2.5.

*Let A*\* *be a solution of* (7). *For all i* = 1, …, *N and all t* ∈ [0, *T* ] *we have* 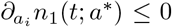 *and* 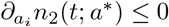, *and therefore* 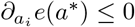.

**Proof**. Note that as *n*_1_ and *n*_2_ are differentiable in *W* ^1,∞^(), *T*) implies that they are differentiable in *C*^0^([0, *T* ]). Using the boundedness of *n*_1_ and *n*_2_, see Proposition 2.1, we get

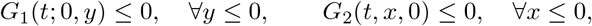

which in combination with (15)-(18) implies 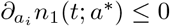 and 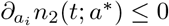, and therefore 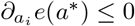, which proves the lemma. □

#### Remark 2.6.

*This lemma suggests that the solution a*\* *should be on the boundary of* 𝒜. *The following theorem shows that indeed A*\* ∈ *∂*𝒜 *and typically a*\* *is a bang-bang type*.

#### Theorem 2.7.

*Assume* 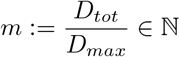. *Let* 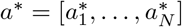 *be a solution of* (7) *and*

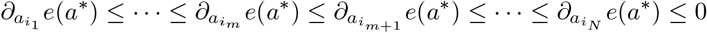

*be an increasing ordering of components of* 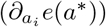. *We have*

i. *If* 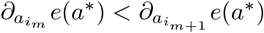 *then the solution a*\* *is of bang-bang type. More precisely*

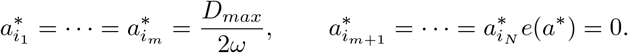
ii. *In general, if*

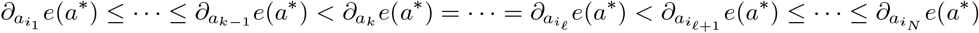

*for certain k*, 𝓁 *integers such that* 1 ≤ *k* ≤ *m* < 𝓁 ≤ *N, then*

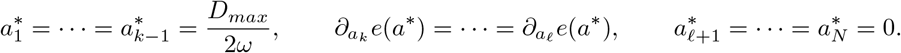

**Proof**. We note that from Lemma 2.5, 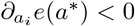 for all 1≤ *i* ≤ *m* in the case i) and 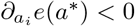 for all 1 ≤*i* ≤ *l* in the case ii).

Let us first prove i). We assume that the claim does not hold. Then necessarily there exists: i.1) an integer *i* with 1 ≤ *i* ≤ *m* and 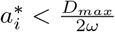, or i.2) an integer *j* with *m* < *j* ≤ *N* such that 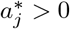.

In the case i.1), if all 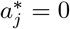,*m* < *j* ≤ *N*, we consider *z* = (0, …, 0, 1, 0, …, 0), where 1 is on *i*the place and *a* = *a*\* + *hz, h >* 0 small. Then *a* ∈ 𝒜 and

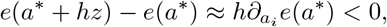

which is a contradiction and proves that this case cannot happen. Otherwise, if 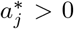 for any integer *j, m* < *j* ≤ *N*, we consider *z* = (0, …, 0, 1, 0, …, 0, −1, 0, …, 0), where 1 is on *i*the place and −1 in *j*th place. Then *a* = *a*\* + *hz* ∈ 𝒜 for *h >* 0 small and

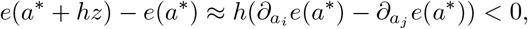

which is again a contradiction and shows that this case does not happens.

In the case i.2), from the constraint (5) necessarily there exists an integer *i*, 1 ≤*i* ≤ *m* with 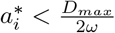. Then we consider *z* = (0, …, 0, 1, 0, …, 0, −1, 0, …, 0), where 1 is on *i*the place and 1 in *j*th place and conclude as in the case i.2) above.

For the proof of ii), if *k* = 1 and 𝓁 = *N*, there is nothing to prove - the vector 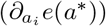 is constant. The cases *k >* 1 or 𝓁 < *N* are proved similarly as i) above. Indeed, if we assume that the claim does not hold then there exits: ii.1) an integer *i*, 1 ≤ *i* ≤ *k*, with 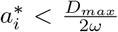, or ii.2) an integer *j*, 𝓁 + 1 ≤ *j* ≤ *N*, with 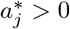.

In the case ii.1), resp. ii.2), we proceed as in i.1), resp. i.2), above and conclude that this case cannot happen. □

#### Remark 2.8.

*In all our numerical experiments we have found optimal solution a*\* *only of the type i) of Theorem 2.7. We conjecture that the case ii) of Theorem 2.7 does not happen*.

### 2.3 Numerical optimization

To solve (7), we can use a gradient method. Note that our minimization problem is constrained and the set 𝒜 is convex. To satisfy the constraints for *a*, the constrained optimization problem (7) is solved with the projected gradient descent method (see [13]) as given by Algorithm 1. We note that *a*^*k*+1^ of the problem (22) below could also be solved by using a quadratic programming algorithm,

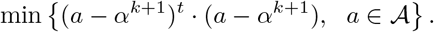

#### Algorithm 1: Gradient descent method

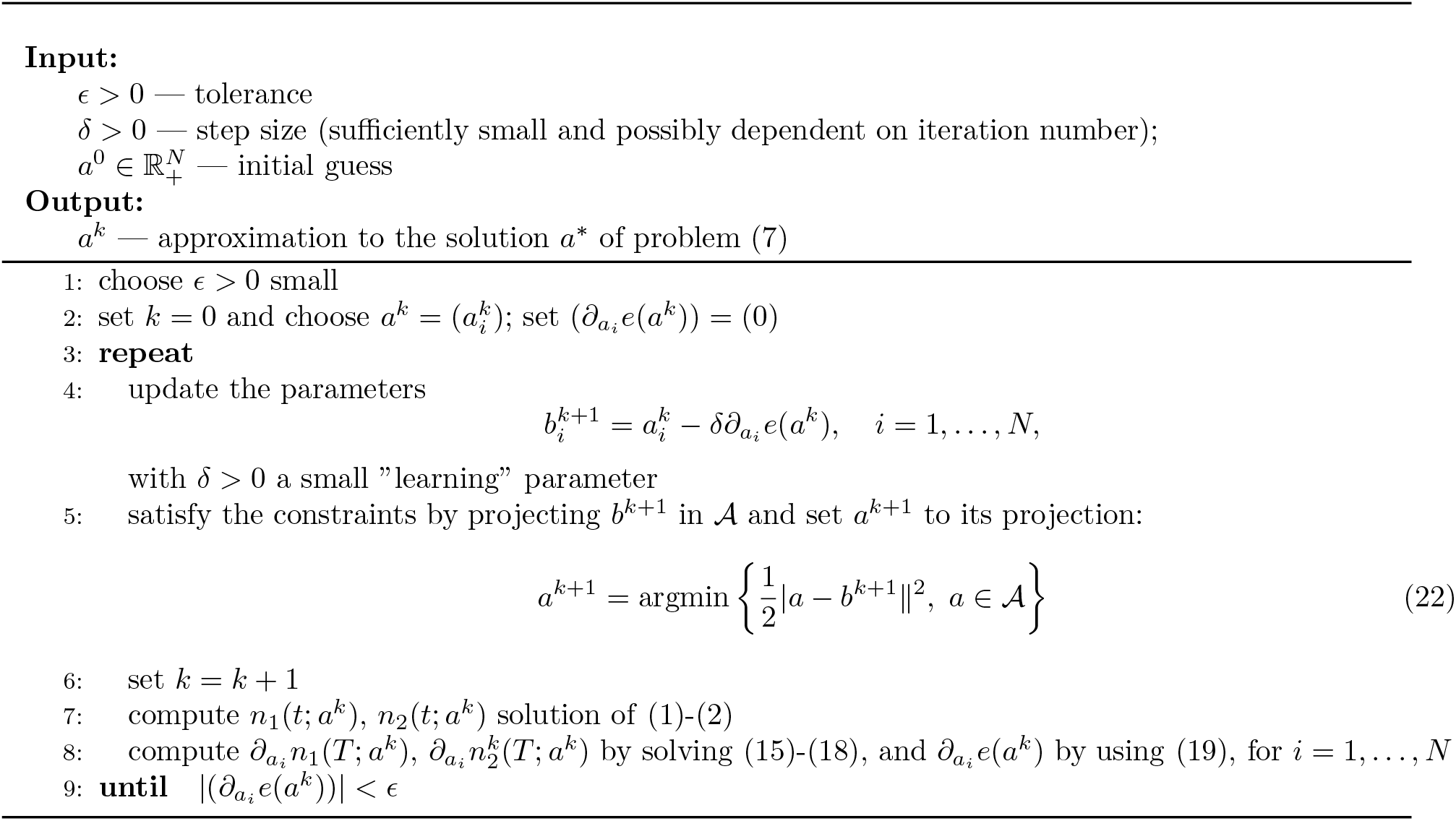

Computing (19) is expensive because we have to solve (15)-(18) for all *i* = 1, …, *N*. We can reduce the computational cost by using appropriate adjoint functions as follows.

#### Proposition 2.9.

*Let a* ∈ ℝ^*N*^. *Then the system of ODEs*

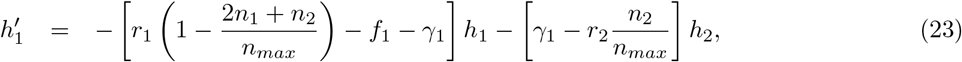

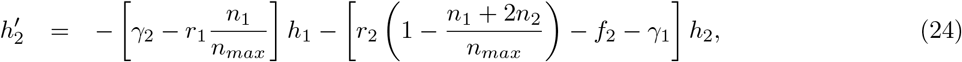

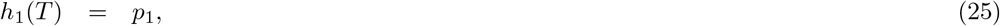

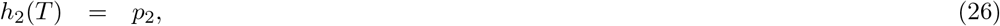

*has a unique solution* (*h*_1_, *h*_2_) ∈ *W* ^1,∞^(((0, *T*); ℝ^2^). *Furthermore*,

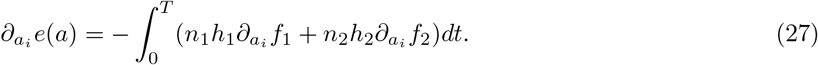

**Proof**. By changing the variable *s* = *T* − *s, s* ∈ (0, *T*) and setting *g*_*i*_(*s*) = *h*_*i*_(*T* − *s*), *i* = 1, 2, we find that (*g*_1_, *g*_2_) solves a system of (linear) ODEs similar to (23)-(26) but with initial conditions at *s* = 0. Then we prove the existence and solution of (*g*_1_, *g*_2_), and so of (*h*_1_, *h*_2_), by proceeding as in Theorem 2.2 for (*g*_1_, *g*_2_) instead of (*n*_1_, *n*_2_).

Now we prove formula (27). Note that 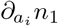 and 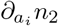 solve (15)-(18). Then

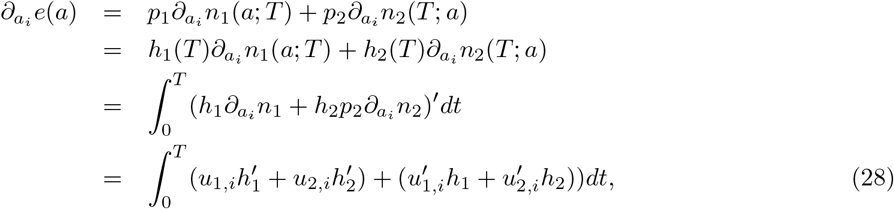

where for simplicity we wrote 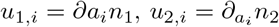. Using (15)-(18) we can evaluate 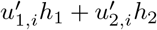 as follows

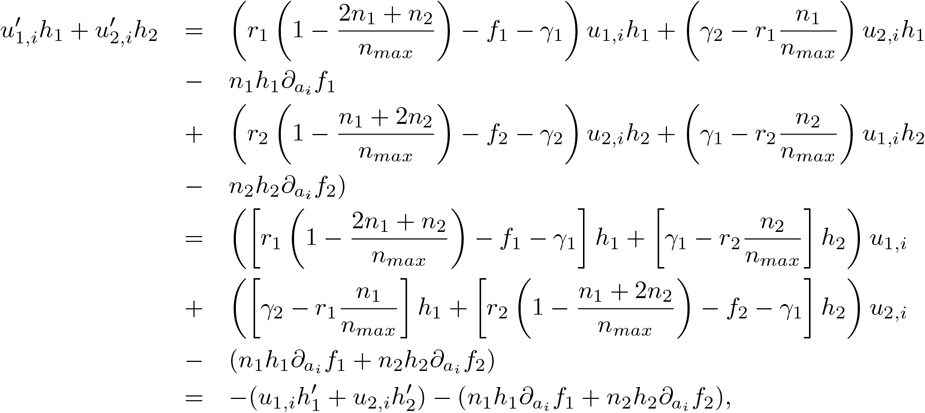

which in combination with (28) proves (27).□

Based on (27) we amend the Algorithm 1 as follows:

#### Algorithm 2: Gradient descent method amended

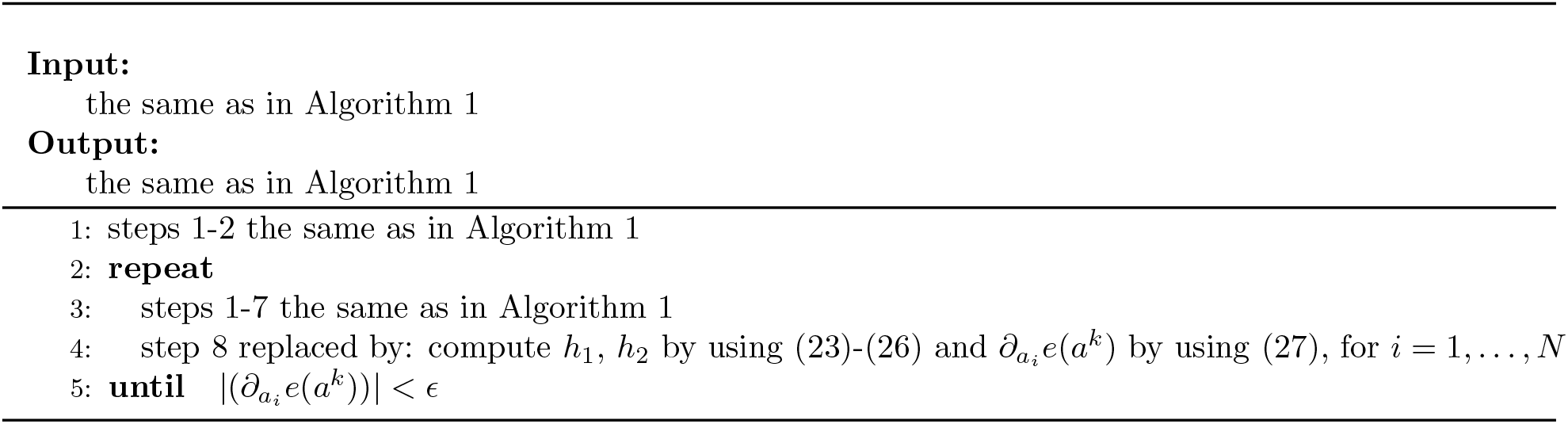

This optimization procedure is implemented in python. To solve the system of ODEs, the `lsoda` method is used, and to compute the projected gradient, the `solve_ls` function from `qpsolvers` is used.

## 3 Results

As the set of variables to consider is large, we make several assumptions on the radiation schedule before attempting optimization. We choose *N* = 30, *D*_*max*_ = 3Gy, and *D*_*tot*_ = 60Gy and consider the end of the treatment period to be the end of the 31^*st*^ day. This choice is easily changeable, but fixing it allows us to perform analysis. Since *ND*_*max*_ *> D*_*tot*_, the optimal solution cannot simply be to apply maximum radiation at every fraction. Additionally, since a nondimensionalization of cell number would eliminate the parameter *n*_*max*_, we set *n*_*max*_ = 1 for convenience - the results of the model are independent of the choice of *n*_*max*_.

Additionally, tumour cells typically have a known value for their ratio of radiobiological parameters, the *α/β* ratio. This ratio is frequently cited as being in the range 3-10 *Gy*, and fixing it allows us to greatly speed up numerical calculations. Furthermore, the effects of this ratio have been extensively studied in other works, and our primary purpose is to investigate the effect of radiation altogether, rather than the effect of each part of the LQ model. For these reasons, we fix *α/β* = 3 for all simulations. We also assume that prior to radiation, the nondimensionalized cell fraction has reached its steady state. This steady state is easily obtained from the governing ODEs by setting the derivatives and the terms with *f*_1_, *f*_2_ to zero and solving for the cell numbers as *n*_1_(0) = 1*/*(1 + *γ*_1_*/γ*_2_) and *n*_2_(0) = 1*/*(1 + *γ*_2_*/γ*_1_), which we use as our initial condition. Using the optimization procedure outlined above, the optimal distribution of radiation can be identified for any given parameter set which consists of the eight parameters found in equations (1), (2), (3), and (4) (not including *n*_*max*_). In order to complete the optimization, we must also specify the metric function weight coefficients, *p*_1_ and *p*_2_.

When using the optimization procedure on various parameter sets, a dominant qualitative behaviour quickly emerges; specifically, that the optimal radiation schedule is when the final 20 days receive the maximum allowable radiation and the first 10 days receive none. We refer to this case as the ‘trivial’ result since it is both the least interesting and most common among reasonable parameter sets. Intuitively, the reason for this result is that since we compute the metric at the end of treatment, the optimization tends to find that focusing radiation at the end of treatment does not allow time for regrowth. However many parameter sets do not fall into this pattern and show more varied optimal dose distributions. We refer to cases which do not optimize to the trivial result as ‘nontrivial’ results. For example, consider the optimization results shown in Figure 2. In this figure, we seek to minimize the total cell number and therefore choose *p*_1_ = *p*_2_ = 1. We generate the parameter set of *r*_1_ = 0.4, *r*_2_ = 0, *α*_1_ = 0.2, *β*_1_ = *α*_1_*/*3, *α*_2_ = 0.0005, *β*_2_ = *α*_2_*/*3, *γ*_1_ = 0.1, *γ*_2_ = 0.1 and use our procedure to find the radiation distribution that minimizes our metric. Notice that the optimal distribution of radiation is to apply the maximal amount of radiation on all days other than days 3, 4, 6, 9, 10, 13, 16, 20, 24, and 27. In fact, this schedule results in an approximate 22% decrease in final cell number compared to the clinical case and an approximate 40% decrease in final cell number compared to the trivial case.

**Figure 2:**
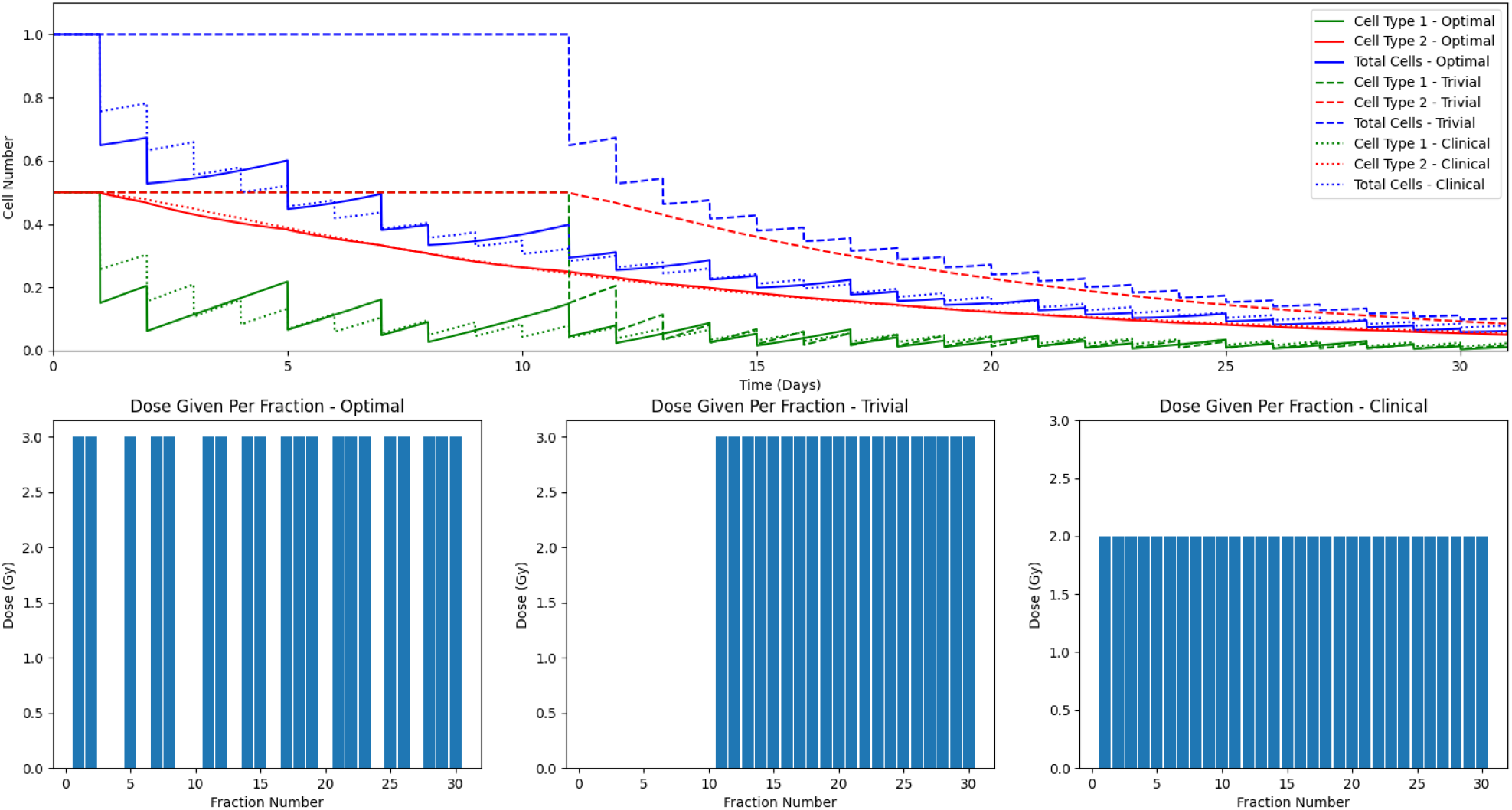
A comparison of model results with different radiation schedules. Cell type 1 is shown in green, cell type 2 in red, and total cells in blue. The schedule derived using the optimization procedure is plotted with the full line, the trivial case with the dashed line, and the clinical standard case with the dotted line. The parameters used for this case are: *r*_1_ = 0.4 (1/day), *r*_2_ = 0 (1/day), *α*_1_ = 0.2 (1/Gy), *β*_1_ = *α*_1_*/*3 (1/Gy), *α*_2_ = 0.0005 (1/Gy), *β*_2_ = *α*_2_*/*3 (1/Gy), *γ*_1_ = 0.1 (1/day), *γ*_2_ = 0.1 (1/day). Notice that the optimal case results in an approximate 22% decrease in final cell number compared to the clinical case and an approximate 40% decrease in final cell number compared to the trivial case.

Cases such as the one displayed in Figure 2 are the outliers however, with the trivial distribution being the optimal for most parameter sets. This then begs the question of which parameter sets result in nontrivial optima and what are the key parameters which determine such behaviour. Answering these questions using analytical methods is quite challenging due to the number of variables included in the model and the complexity of the optimization process. We therefore result to numerical and data analysis techniques to understand the relationship between the parameters and the optimal radiation distribution.

### 3.1 Minimization of total cells

We seek to identify the relationship between the model parameters and the optimal radiation schedule for the case that the clinical goal is to kill the maximal number of cells. For this, we set *p*_1_ = *p*_2_ = 1 for our optimization and attempt to identify the subset of the parameter space which produces nontrivial optimization results. Due to the large number of model parameters and complex nature of the optimization algorithm, it is challenging to use analytical tools to partition the parameter space. So instead, we use numerical and data analysis techniques to derive qualitative insights.

We begin by generating an input-output data set where each sample consists of the values for the six free parameters along with a label of trivial or nontrivial for its optimization result. To create this dataset, we select clinically-reasonable ranges for each of the parameters and sample values from within those ranges. The ranges used for sampling from the parameter space are shown in Table 1. Note that for the proliferation and plasticity rates, parameters are uniformly sampled, whereas for the radiobiological parameters, they are sampled on a logarithmic scale. As a full optimization typically takes several minutes to find the optimum, in order to generate a large enough dataset for analysis, we slightly alter the optimization. Since we are only interested in labelling the sample as trivial or nontrivial, we do not need to complete the full optimization process. Instead, we can label the sample by setting our initial optimization guess as the trivial result, performing a single optimization iteration, and determining whether or not the new optimal guess has changed: if it has, then the trivial case must not be the optimal solution and there must therefore be a nontrivial result. This method allows for the generation of a much larger dataset than would otherwise be possible given computational limitations. With this method, a data set of 19,338 labelled parameter samples are generated and used for analysis

**Table 1:** Parameter sampling ranges and method for the minimization of total cells case.

| Parameter | Symbol | Range Min (Unit) | Range Max (Unit) | Sample Type |
| --- | --- | --- | --- | --- |
| Cell Proliferation Rate | $r_i$ | 0.3466 (1/day) | 0.0347 (1/day) | Uniform |
| Linear LQ Parameter | $\alpha_i$ | $e^{-10}$ (1/Gy) | 1 (1/Gy) | Logarithmic |
| Cell Plasticity Rate | $\gamma_i$ | 0 (1/day) | 0.3 (1/day) | Uniform |

To identify which parameters are the most important in determining the optimization result, we create a 2D histogram matrix as can be seen in Figure 3. In the upper triangular portion of this figure, a 2D histogram of each pair of model parameters is plotted with blue representing trivial results and orange representing nontrivial results. As can be clearly seen from these histograms, a separation between the optimization result is most apparent when considering the effect of radiation through the parameters *α*_1_ and *α*_2_ (plotted in this Figure on a log scale). Interestingly, the *α*_1_ vs. *α*_2_ histogram shows that nontrivial results are grouped along the axes, where one of the values is relatively large and the other relatively small. This motivates the idea that a key determining factor in the result is the ratio of the radiation effect parameters between the cell types. In the bottom part of Figure 3, the 2D histogram of the logarithm of the larger *α*_*i*_ is plotted against the logarithm of the larger *α*_*i*_*/α*_*j*_. A clear separation between the results is observed in this plot, showing that the most important factor is the difference in radiation effect between the cell types. To further illustrate this, a tree classifier (Figure 4) was created and trained using only *α*_1_ and the ratio *α*_1_*/α*_2_. Limited to only this knowledge and a maximum decision depth of three, the tree classifier was able to obtain 88.6% accuracy. The tree classifier was created using the `DecisionTreeClassifier()` function from `sklearn`

**Figure 3:**
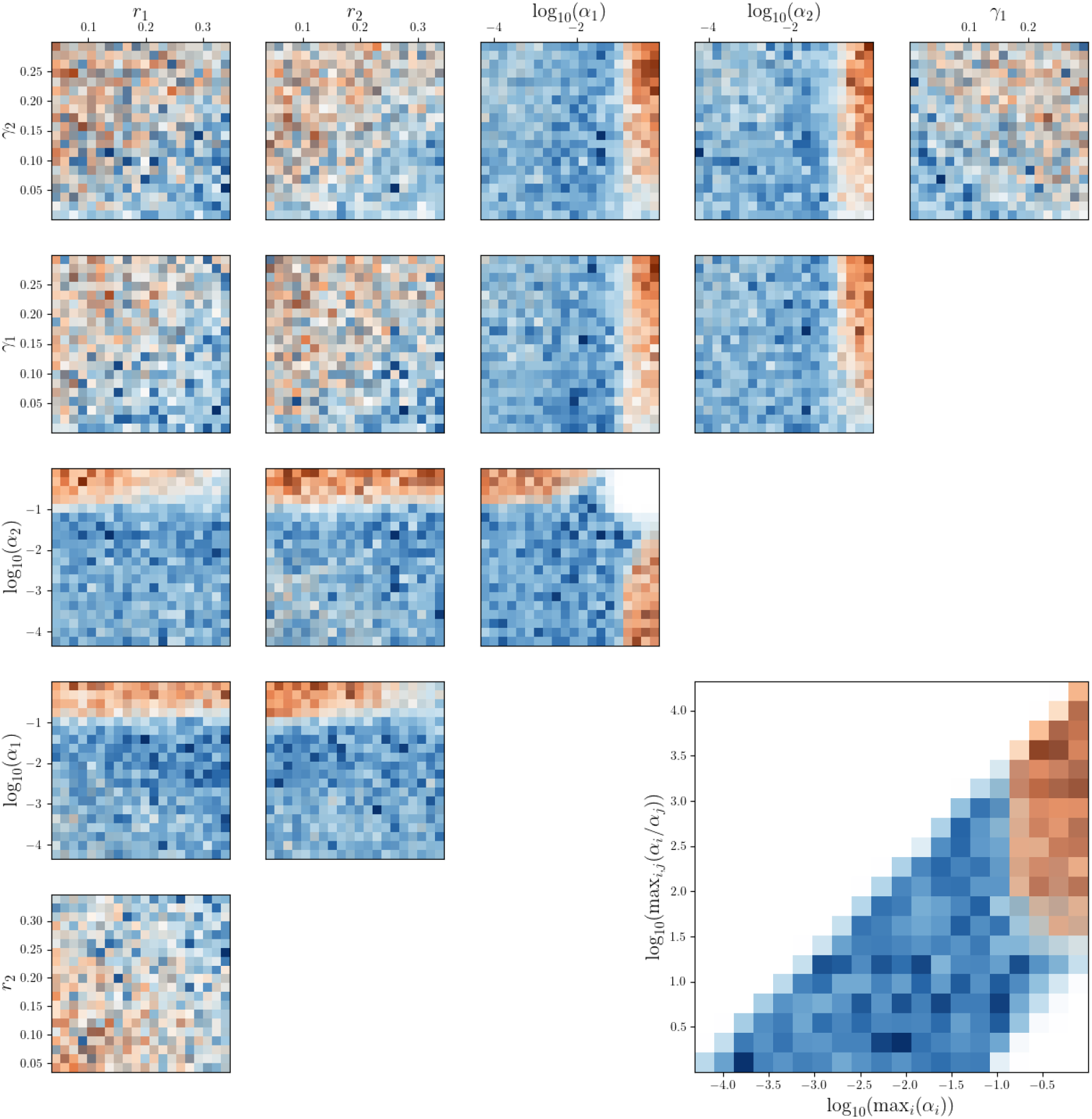
A 2D histogram matrix of each pair of model parameters for the case of minimization of total cells. Areas that are more solid blue represent cases where the optimal result was the trivial case and areas of more solid orange represent cases where the optimal result was nontrivial. Note that the parameters *α*_1_ and *α*_2_ are plotted on a base 10 log scale. Observe the clear separation in cases involving *α*_1_ and *α*_2_ and the lack of separation in cases that don’t. In the bottom portion of the figure, a 2D histogram with the same colour scheme is included for the maximum *α*_*i*_ vs. the maximum *α*_*i*_*/α*_*j*_, with both axes on a base 10 log scale. Notice the clear separation between the optimization cases based on the ratio of the radiation effect parameters. The parameters *r*_1_, *r*_2_, *γ*_1_, and *γ*_2_ are in units of (1/day) while *α*_1_ and *α*_2_ are in units of (1/Gy).

**Figure 4:**
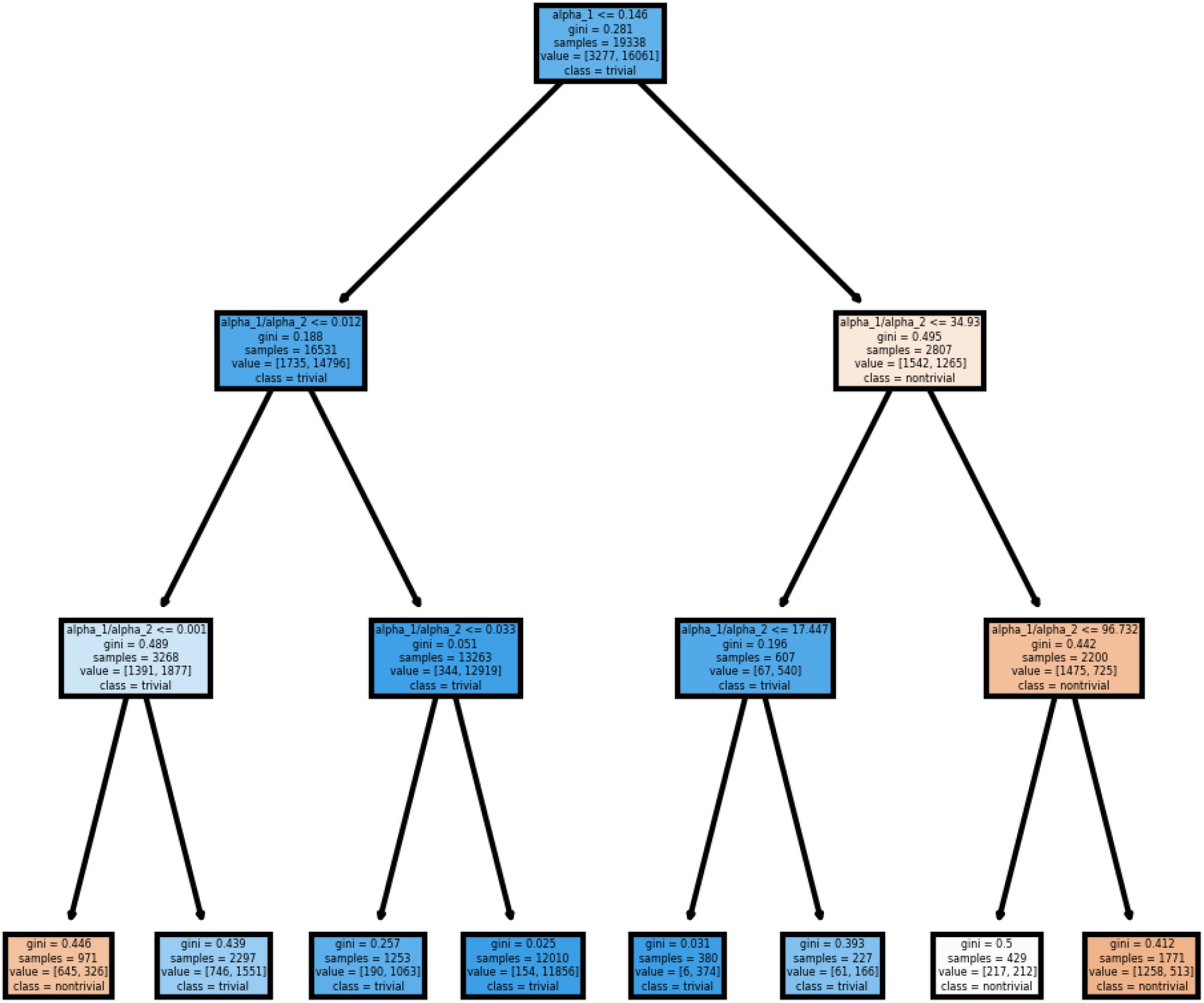
A tree classifier for the minimization of total cells case. The tree uses only *α*_1_ and the ratio *α*_1_*/α*_2_ to make predictions and is limited to a maximum decision depth of three. Even with these limitations, the tree is nonetheless able to obtain 88.6% accuracy, showing that the qualitative nature of the optimization result is largely determined by the radiation effect parameters.

In light of these results, it is worth noting that the situation where two distinct populations of cells have differing levels of radiation response is precisely what has been observed in previous studies examining CSC subpopulations. Specifically, [15] and [21] both observed that CSCs exhibit less sensitivity to radiotherapy than non-CSCs. Furthermore, revisiting the disagreement between the results of Leder et al [16] and Forouzannia et al [11] compared with Galochkina et al [12], we can more concretely evaluate our hypothesis that the reason for the disagreement was due to the different assumptions about the effect of radiation on the cell types. We find that in cases where the effect of radiation is constant between the cell types, the optimal schedule of radiation is trivial. With the increasing ability of researchers to identify CSC subpopulations within tumours, results such as these are clearly of importance for treatment planning.

### 3.2 Minimization of cancer stem cells

Given the importance of CSCs to the progression of cancer, it has been proposed that treatments should prioritize the elimination of CSCs rather than the minimization of total cells. Accordingly, we consider the case where *p*_1_ = 0 and *p*_2_ = 1 to see the different strategies that evolve when seeking to minimize the size of a specific subpopulation. To do so, we designate *n*_2_ as the CSCs and *n*_1_ as the non-CSCs and restrict our parameter generation to align with this distinction. Specifically, given the results of the previous case, we generate samples where 0.1 ≤ *α*_1_≤ 1 and 20 < *α*_1_*/α*_2_ < 100 and create another histogram matrix. The results of this are shown in Figure 5 for a set of 17853 samples. In this figure, notice that there is a clear separation between the trivial and nontrivial cases on the plot of *r*_2_ vs. *γ*_2_. Importantly, the case of a tumour subpopulation which exhibits radioresistance, a small growth rate, but a high replenishment rate of normal cancer cells is precisely the behaviour expected of CSCs. These results buttress our qualitative conclusion from the last section that non-uniform radiation schedules can be preferable in cases where there is a known CSC population.

**Figure 5:**
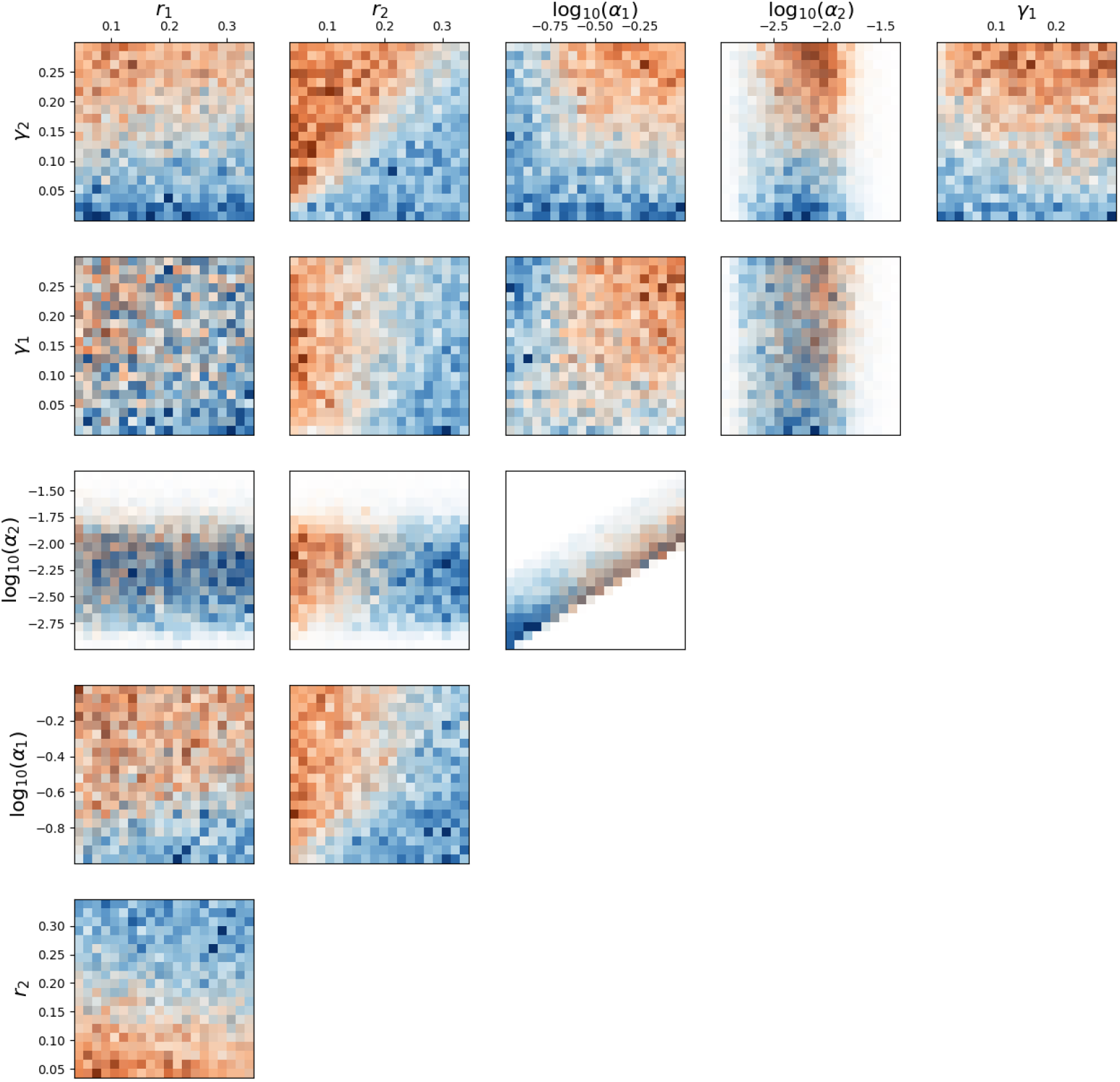
A 2D histogram matrix of each pair of model parameters for the case of minimization of CSCs. Areas that are more solid blue represent cases where the optimal result was the trivial case and areas of more solid orange represent cases where the optimal result was nontrivial. Note that the parameters *α*_1_ and *α*_2_ are plotted on a base 10 log scale. Observe that for minimization of CSCs with a restricted parameter space, the clearest separation is now seen in the plot of *r*_2_ vs. *γ*_2_. The parameters *r*_1_, *r*_2_, *γ*_1_, and *γ*_2_ are in units of (1/day) while *α*_1_ and *α*_2_ are in units of (1/Gy).

## 4 Conclusion

In this paper, we added a crucial element to the classic problem of temporal optimization of radiation – cellular heterogeneity. We created a differential equation model and an optimization procedure to identify which distribution of radiation dose resulted in the maximal cell kill for a given set of parameters governing a tumour’s growth and response to treatment. We found that most simulated sets of governing parameters resulted in the optimal distribution being when the allowable dose was focused as much as possible at the end of treatment, a case which we termed the trivial result. But some parameter sets exhibited nontrivial results where the available dose was spread across the treatment time. Of note, we showed that the optimal distribution is typically bang-bang, with each day’s radiation either being the maximum allowable or zero. We find that the main factor in determining whether the optimal radiation dose will be nontrivial is the difference in the effect of radiation between the cell types. We specifically showed that knowledge of the linear LQ parameters for the cell types alone was enough to predict the behaviour with over 88% accuracy. There are several parts of our work that could be further generalized and limitations of our model which provide opportunities for future research directions. The assumptions made in creating our model can easily be changed: one could consider a Gompertzian growth, or a nonlinear plasticity, for example. The radiation schedule could also be generalized to include fractions of different lengths and start times. Continuous dose profiles could also be worked into the optimization which may be more appropriate in cases such as when the method of administration is brachytherapy. The assumptions made on our choices of parameters could also be changed or expanded. Different *α/β* ratios are not included in this study since we are mainly interested in overall radiation effect, though it could be included to provide more general results or fixed to a value for a particular patient for more specific recommendations. These results could also be taken further, and one could attempt to find the precise relationship between the parameters and the qualitative behaviour beyond just the difference in radiation effect, though this would likely require a more efficient optimization procedure or significantly more computing power. Furthermore, actually predicting which days would receive radiation rather than just the qualitative behaviour would be interesting, though again, this would certainly require more data or computing power.

Our results shed important light on the question of how differences in tumour subpopulations can lead to changes in optimal treatment protocol. This is particularly relevant when discussing CSCs, which are known to exist in tumours and display radioresistance compared to non-CSCs. We hope that this work acts as support for the idea that understanding the full tumour biology is important for designing effective treatments and the idea that mathematical tools can play a vital role in identifying cases where treatments can be improved. We hope that the results presented here can be used as hypotheses for clinical and experimental researchers to test our findings. Clearly, the recommendations which could arise from our results are of importance to clinical practice.

## Notes

### Competing Interest Statement

The authors have declared no competing interest.

